# Odorant cues linked to social immunity induce lateralized antenna stimulation in honey bees (*Apis mellifera L*.)

**DOI:** 10.1101/086330

**Authors:** Alison McAfee, Troy F. Collins, Lufiani L. Madilao, Leonard J. Foster

## Abstract

Hygienic behaviour (HB) is a social immunity trait in honey bees (*Apis mellifera* L.) whereby workers detect, uncap and remove unhealthy brood, improving disease resistance in the colony. This is clearly economically valuable; however, the molecular mechanism behind it is not well understood. The freeze-killed brood (FKB) assay is the conventional method of HB selection, so we compared odour profiles of FKB and live brood. Surprisingly, we found that significantly more brood pheromone (β-ocimene) was released from FKB. β-ocimene abundance also positively correlated with HB, suggesting there could be a brood effect contributing to overall hygiene. We found that &#x03B2; ocimene stimulated worker antennae in a dose-dependent manner, with the left antennae responding significantly stronger than right antennae in hygienic bees, but not in non-hygienic bees. This suggests that HB depends not only on odour detection, but also lateralization of sensitivity. We also compared odour profiles of *Varroa-*infested brood to healthy brood and found an overall interactive effect between developmental stage and infestation, but specific odours did not drive these differences. Overall, the data we present here is an important foundation on which to build our understanding the molecular mechanism behind this complex behaviour.

## Introduction

Honey bees (*Apis mellifera* L.) face many challenges, but disease is perhaps the most significant^1^. Of the diseases and parasites that bees are susceptible to, the ectoparasite *Varroa destructor* is the most devastating^2-4^, owing to its rapid spread around the globe and its ability to simultaneously parasitize brood and transmit viral pathogens^4,5^. Although there have been attempts^6,7^, there is currently no commercial treatment for the virus by mites. Measures for controlling *Varroa* do exist, including formic acid, oxalic acid, fluvalinate and coumaphos, among others; however, these treatments also stress the bees themselves, are laborious for the beekeeper and require precise timing of treatment to avoid economic losses^8,9^. Furthermore, some of the acaricides are fat-soluble and accumulate in the beeswax over time, contaminating harvested hive products like wax, propolis and honey^10^. More concerning, however, are reports of chemical resistance in mites^11,12^. For these reasons, there has been considerable interest in breeding bees that are resistant to mites themselves^13-17^.

Honey bees bred for HB have higher survival rates when challenged with *Varroa*, American foulbrood and chalkbrood^13,18,19^. Bees perform HB by detecting, uncapping and removing diseased brood from the colony to reduce pathogen load, thereby improving disease resistance^20,21^. The most well-established method of selecting for hygienic bees is the FKB assay, in which patches of brood are frozen with liquid nitrogen, returned to the hive and evaluated after 24 h. The HB score is defined as the fraction of dead pupae have been detected and removed^20,21^. Colonies that perform well in this test also have improved outcomes when challenged with real diseases, allowing FKB to be an effective tool for selective breeding^20^.

There is a large body of evidence suggesting that hygienic bees identify diseased brood through olfactory cues^15,16,22-26^ and that they are more sensitive to and better at discriminating between them^22,26^. The antennae, bees’ main olfactory organ^27^, have been shown to play a pivotal role in HB with multiple independent research groups identifying significantly differentially expressed antennal genes in hygienic *versus* non-hygienic bees, as well as strong antennal biomarkers for selective breeding^14-17,28^. Odorant binding protein (OBPs) aid odour detection and are consistently upregulated in hygienic bees’ antennae. However, relatively little is known about precisely what odours the bees are detecting. One study investigated the volatile odours emitted from chalkbrood-infected larvae^25^ and several focused on possible cues from *Varroa*-infested brood^29-33^, but none investigated how they compare to FKB (the main selective test for HB) and very few confirmed the biological activity of the compounds^25,29^. Furthermore, how infested brood odour profiles change with respect to pupa development (and associated growth of the mite family) is yet unknown. In this study, we use gas chromatography-coupled mass spectrometry (GC-MS) to find striking differences in compounds emitted from FKB and *Varroa*-infested brood across developmental stages. We also functionally validate the biological activity of candidate HB-inducing compounds by quantifying the strength with which they stimulate antennae of hygienic and non-hygienic bees, ultimately suggesting that brood pheromones and lateralization of olfactory sensitivity may be key features of HB.

## Results

### FKB-specific compounds

The FKB assay is thought to work equally well using any age of capped brood^20^, so we reasoned that candidate HB-inducing compounds should have consistently high abundances in dead relative to live brood across ages. To test this, we used GC-MS to compare the cuticle molecular profiles of 12 to 17 d old pupae at 1 to 2 d intervals (Fig. 1A). We found that indeed there were strong differences between dead and live brood (three-factor ANOVA; P < 0.000001; F = 597), which interacted significantly with developmental stage (P = 0.0000024; F = 9.72) and compound identity (P < 0.000001; F = 10.7). Young (12 to 15 d old) FKB tended to have more differentially emitted compounds compared to old (16 to 18 d) FKB (Fig. 2). While most of these compounds were age-specific, one compound, oleic acid, was consistently different across all ages. The identity of this compound was confirmed against a synthetic standard (**Table 1; Appendix S1).**

**Figure 1.**
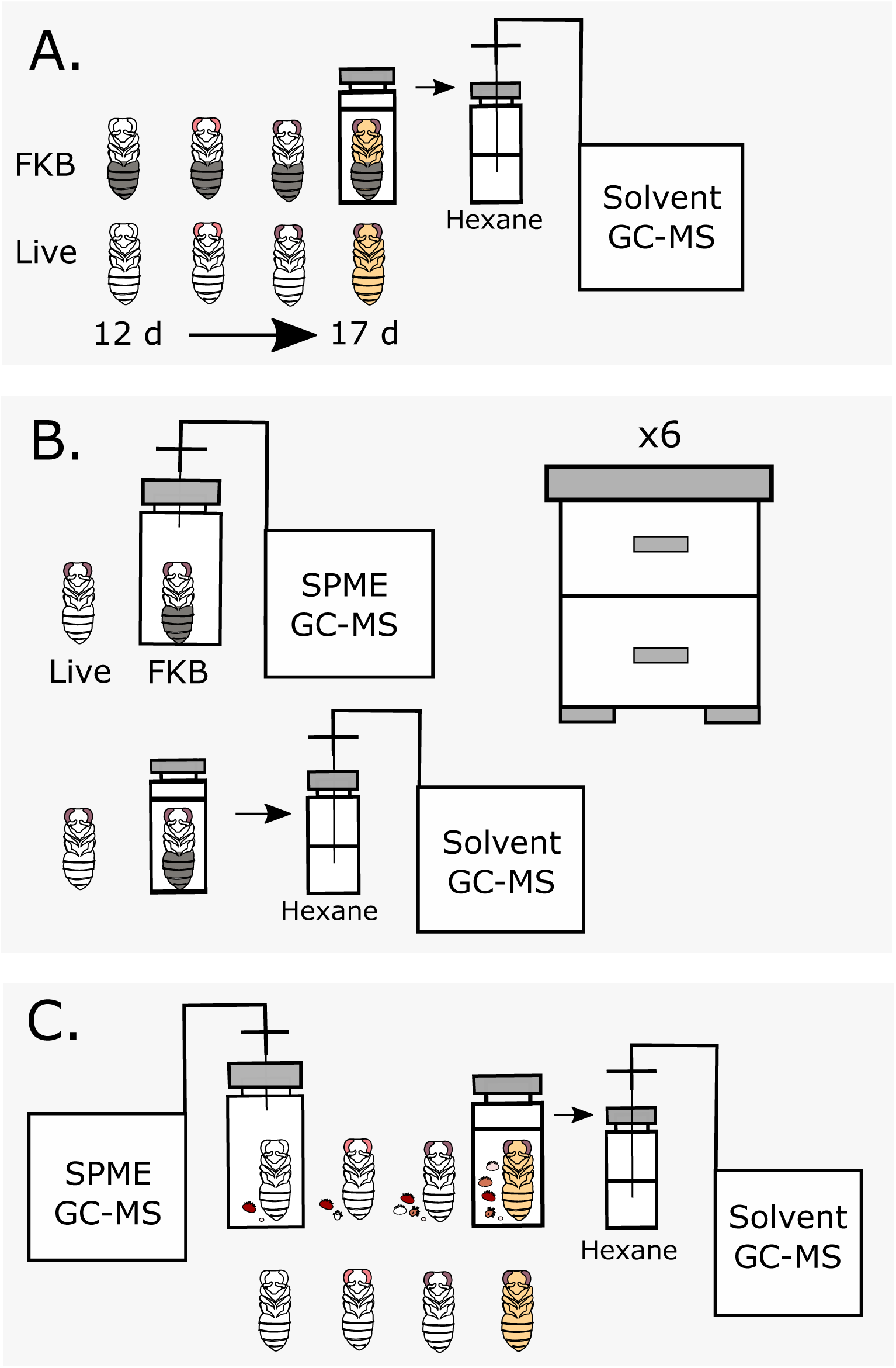
Experimental design schematics. A) Cuticular hydrocarbon analysis. N = 3 for each developmental stage (white-eyed, pink-eyed, purple-eyed white body, purple-eyed tan body). FKB: Freeze-killed brood; GC-MS: gas chromatography mass spectrometry. B) Cross-colony comparison of headspace volatiles and cuticular hydrocarbons. N = 3 for each colony. SPME: solid phase microextraction. C) *Varroa*-infested brood headspace volatiles and cuticular hydrocarbons. Mites and their families were included in each sample. N = 3.

**Figure 2.**
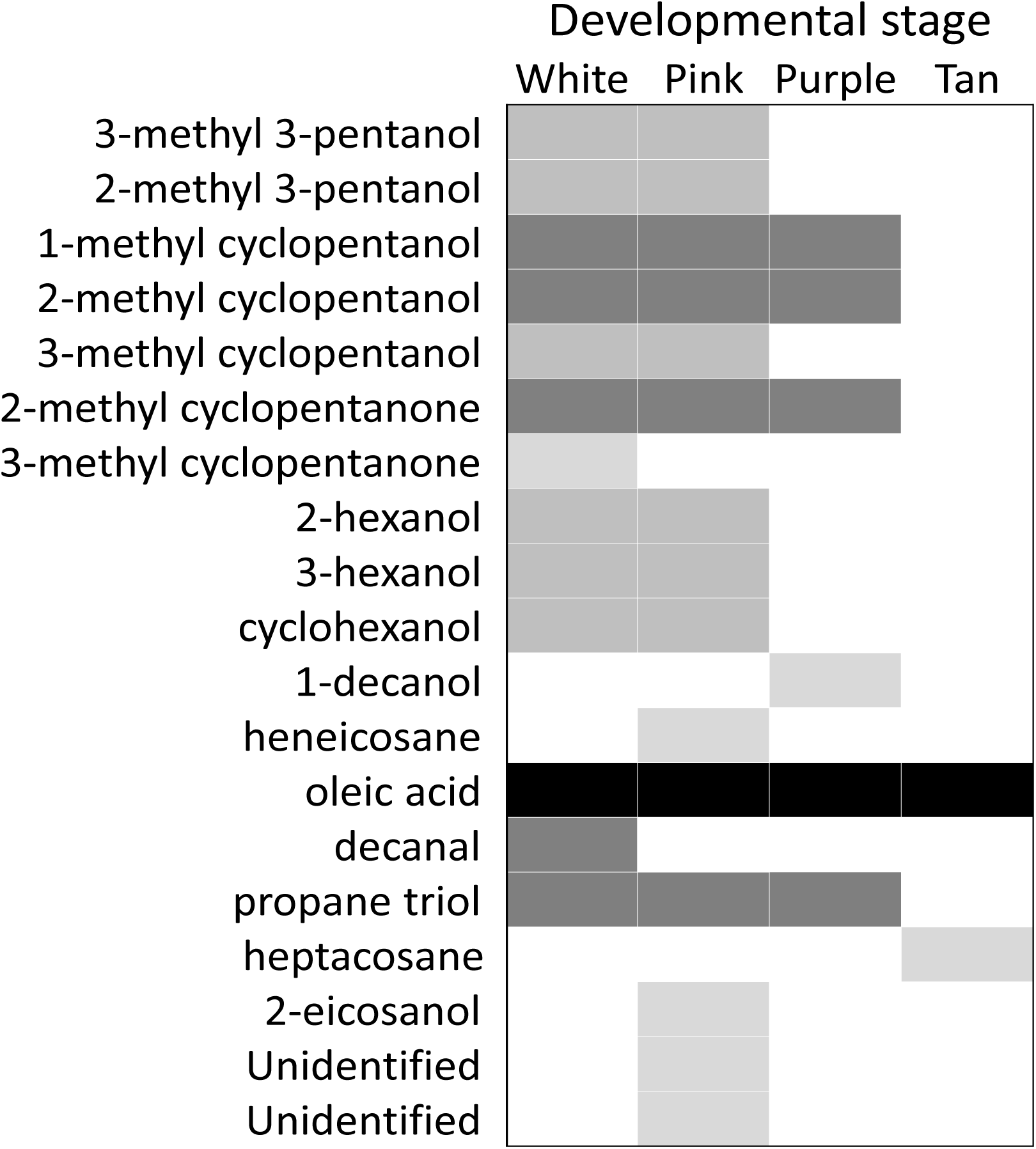
FKB-specific odour profiles vary across developmental stages. Cuticle compounds from live and freeze-killed white-eyed (12-13 d), pink-eyed (14-15 d), purple-eyed white body (16 d) and purple-eyed tan body (17-18 d) were analyzed using gas chromatography mass spectrometry (N=3). Shaded boxes indicate compounds which had significantly different abundances in FKB compared to age-matched live brood. Compounds were identified by comparing mass spectra against a compound library and were not compared to synthetic standards.

We hypothesized that the compounds most likely to be HB-inducers should also be consistently differentially emitted from dead brood across diverse colonies. We compared the odour profiles of FKB to age-matched healthy pupae across six colonies located at three different apiaries. We found ten compounds that were consistently different between FKB and healthy pupae (Fig. 3A and 3B), although the identities of only four (isopropanol, 2-pentanone, β-ocimene and oleic acid) could be confirmed with synthetic standards (**Table 1; Appendix S1**). For five unknowns (Compounds 1 to 5), either the retention times of the synthetic standards did not match the peaks in the samples, making the identifications assigned by the spectral search algorithm unlikely, or the spectra could not be confidently matched to any in the comprehensive Wiley/NIST compound library. Of the ten compounds, nine were most abundant in the FKB headspace samples and only one was most abundant in live pupae. This peak had the highest volatility and a strong 44^+^ base-peak ion, which matches carbon dioxide and is consistent with active respiration.

**Figure 3.**
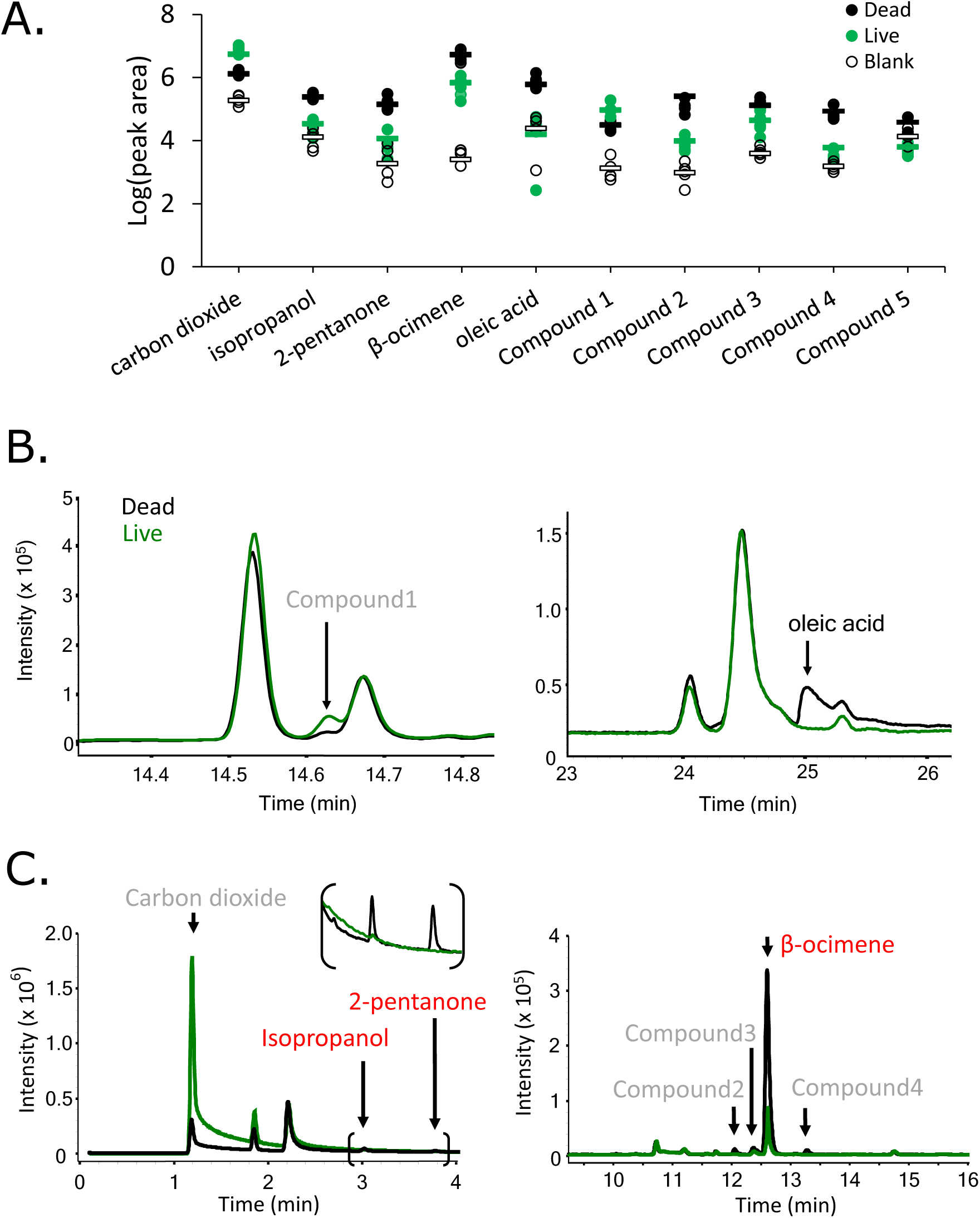
Cross-colony comparison of FKB and healthy brood odour profiles. A total of ten compounds were significantly differentially expressed across colonies (n = 6; two-factor ANOVA; Tukey HSD). A) Solid phase micro-extraction (SPME) method. Bracketed region is enlarged for clarity. Compounds 1 to 4 were identified as 2-methyl tetradecane, α-thujene, α-pinene and 2,3-butanediol, respectively. Compound 5 (not displayed on chromatograms) could not be confidently matched to any spectra in the compound library. Bars represent averages. B) Cuticle hexane wash and C) SPME example chromatograms covering the differentially emitted compounds.

### FKB odour strength is correlated with HB score

It has been established that hygienic adult workers have superior olfactory sensitivity compared to non-hygienic bees^22,23^; however, the brood itself could also play a role in the behaviour^14^. Since a stronger odour should be easier for adult workers to detect and act upon, we hypothesized that brood from highly hygienic colonies may emit a stronger odour signal relative to healthy controls. To test this, we correlated the dead:live ratio of each compound with HB score across eight different colonies. We found that only one compound was significantly correlated with the behaviour: β-ocimene (Fig. 4A; Pearson coefficient = 0.84; P = 0.0059; α = 0.0063; Bonferroni correction). Given that this compound is a familiar brood pheromone that is already known to increase worker visits to cells^34^, this is a remarkable result. Interestingly, β-ocimene was also consistently one of the most intense peaks observed in the chromatograms of dead brood (Fig. 3C), despite – to our knowledge – not being previously thought to be associated with HB.

**Figure 4.**
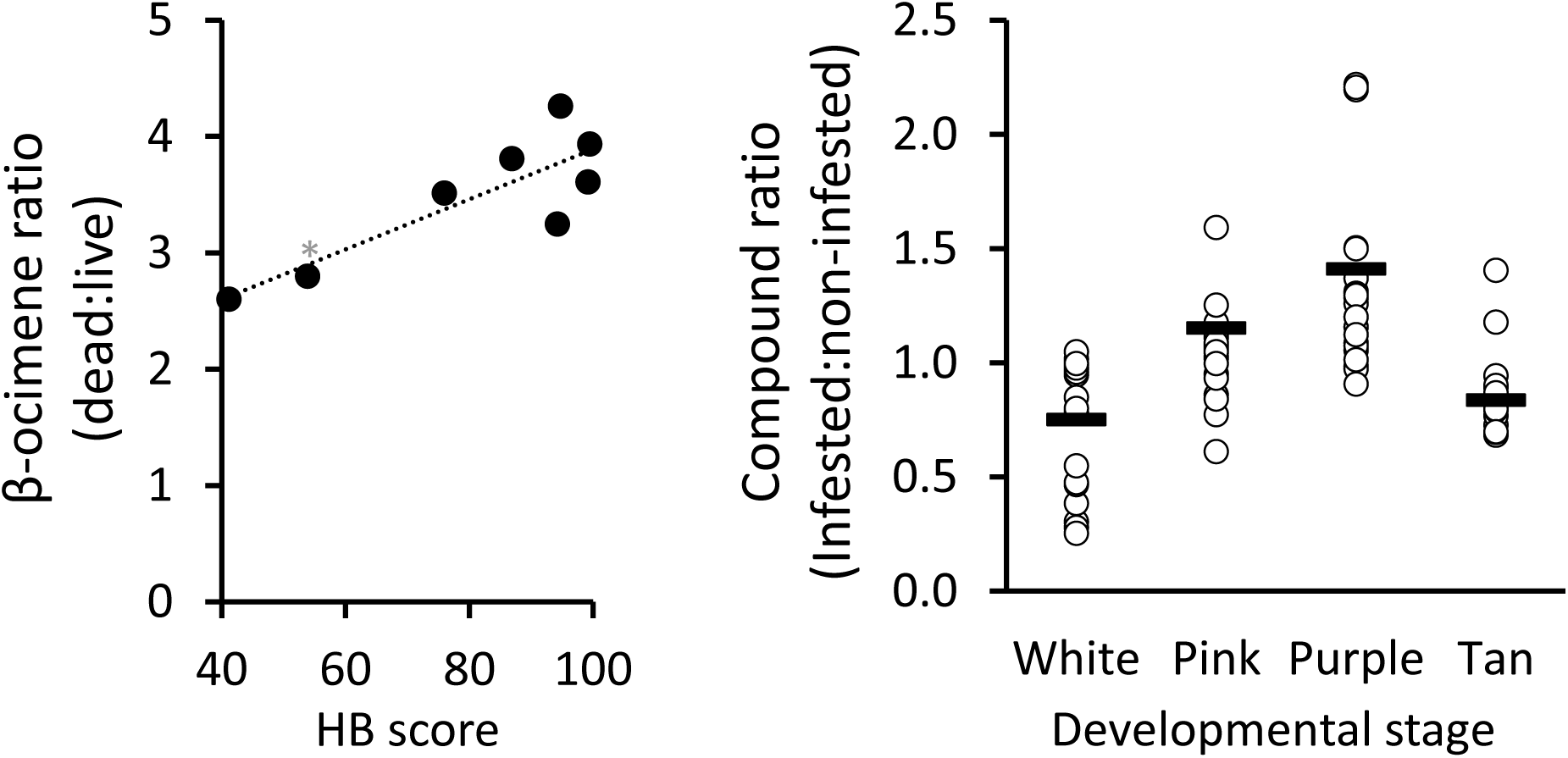
β-ocimene is a key compound in FKB, but not Varroa-infested brood. A) Two rounds of hygienic testing (ref) were performed on eight different colonies. β-ocimene was the only compound to significantly correlate with hygienic behaviour (Pearson correlation coefficient = 0.84, P = 0.0059; N=3 within each colony. *This colony was scored based on one round of hygienic testing. B) *Varroa*-infestation has a significant interacting effect (three-factor ANOVA; P = 0.000022; F = 8.34) on cuticle compound abundance, but this is not driven by specific compounds (P = 0.99; F = 0.38).

### Varroa *infestation interacts with developmental stage to alter cuticle profiles*

To identify chemical cues associated with *Varroa* infestation, we compared odour profiles between infested and non-infested brood. *Varroa* mites reproduce inside the developing pupa’s comb cell, forming a whole family (including the foundress, eggs, protonymphs, deutonymphs and adult males) over time (Fig. 1C). We included four sequential developmental stages (white-eyed, pink-eyed, purple-eyed white body and purple-eyed tan body) and included the mite families with the pupae in the analysis. We did not find any significant effect of infestation in the headspace volatile profile (three-factor ANOVA; P = 0.46; F = 0.56); however, analyzing the cuticle profile showed that while infestation had no effect on its own (three factor ANOVA, P = 0.28, F = 1.15), it significantly interacts with developmental stage (Fig 4B; P = 0.000022; F = 8.34). The overall trend was for infested brood to produce higher levels of compounds relative to healthy brood in age-matched adjacent cells, but no individual compounds drove this effect.

### Electroantennography shows lateralization of olfactory sensitivity in hygienic bees

We investigated the biological activity of isopropanol, 2-pentanone, β-ocimene and oleic acid using electroantennography to quantify antennal nerve depolarizations of hygienic and non-hygienic bees in response to stimuli (Fig. 5). Of all the compounds, only 2-pentanone and β-ocimene showed dose-dependent responses (three-factor ANOVA; P = 0.0002, F = 9.6 and P = 0.0045, F = 5.8, respectively). For β-ocimene, we also found significant interactive effects between dose, level of hygiene (high *vs.* low) and antenna side (left *vs.* right; see Table 1 for P values). Notably, the left antenna of hygienic bees produced the strongest EAG signal overall – significantly higher than the right antennae – whereas non-hygienic bees did not display this effect. This is counterintuitive, since right antennae have a higher proportion of olfactory sensilla^35^ and foragers give stronger EAG responses to (-)-linalool and isoamyl acetate (alarm pheromone) through their right antenna^36^. However, we confirmed that the same left-biased lateralization holds true for another known HB-inducing compound, phenethyl acetate^25^ (isolated from chalkbrood; Fig. 5; 8.5E-10, F = 45.7). Surprisingly, oleic acid appeared not to stimulate bee antennae at all, possibly because of its low volatility at room temperature.

**Table 1.** Differentially emitted compound identifications

| Compound | Base peak<br>(m/z) | Score<sup>1</sup> | Sample<br>RT (min) | Standard<br>RT (min) | Identification<br>accuracy<sup>2</sup> | P-value |
| --- | --- | --- | --- | --- | --- | --- |
| Isopropanol | 44.99 | 90.6 | 3.0 | 3.0 | High | 2.2E-23 |
| 2-pentanone | 43.01 | 91.4 | 3.8 | 3.9 | High | 3.4E-23 |
| *E*-β-ocimene | 93.00 | 94.2 | 12.6 | 12.7 | High | 2.3E-33 |
| Oleic acid | 55.07 | 72.6 | 25.0 | 25.1 | High | 1.8E-70 |
| Compound 1 (α-<br>thujene)<sup>3</sup> | 93.00 | 82.3 | 12.1 | 5.5 | Low | 9.5E-10 |
| Compound 2 (α-<br>pinene) | 93.00 | 94.4 | 12.4 | 5.6 | Low | 2.5E-12 |
| Compound 3 (2,3-butanediol) | 42.98 | 79.2 | 13.3 | 22.4 | Low | 1.2E-13 |
| Compound 4<br>(2-methyl<br>tetradecane) | 57.09 | 83.5 | 14.7 | N/A | Low | 1.6E-45 |

<sup>1</sup>Mass Hunter Qualitative Analysis v.B.06.00
<sup>2</sup>Based on comparison to synthetic standards
<sup>3</sup>Bracketed compound names represent the proposed Mass Hunter matches

**Figure 5.**
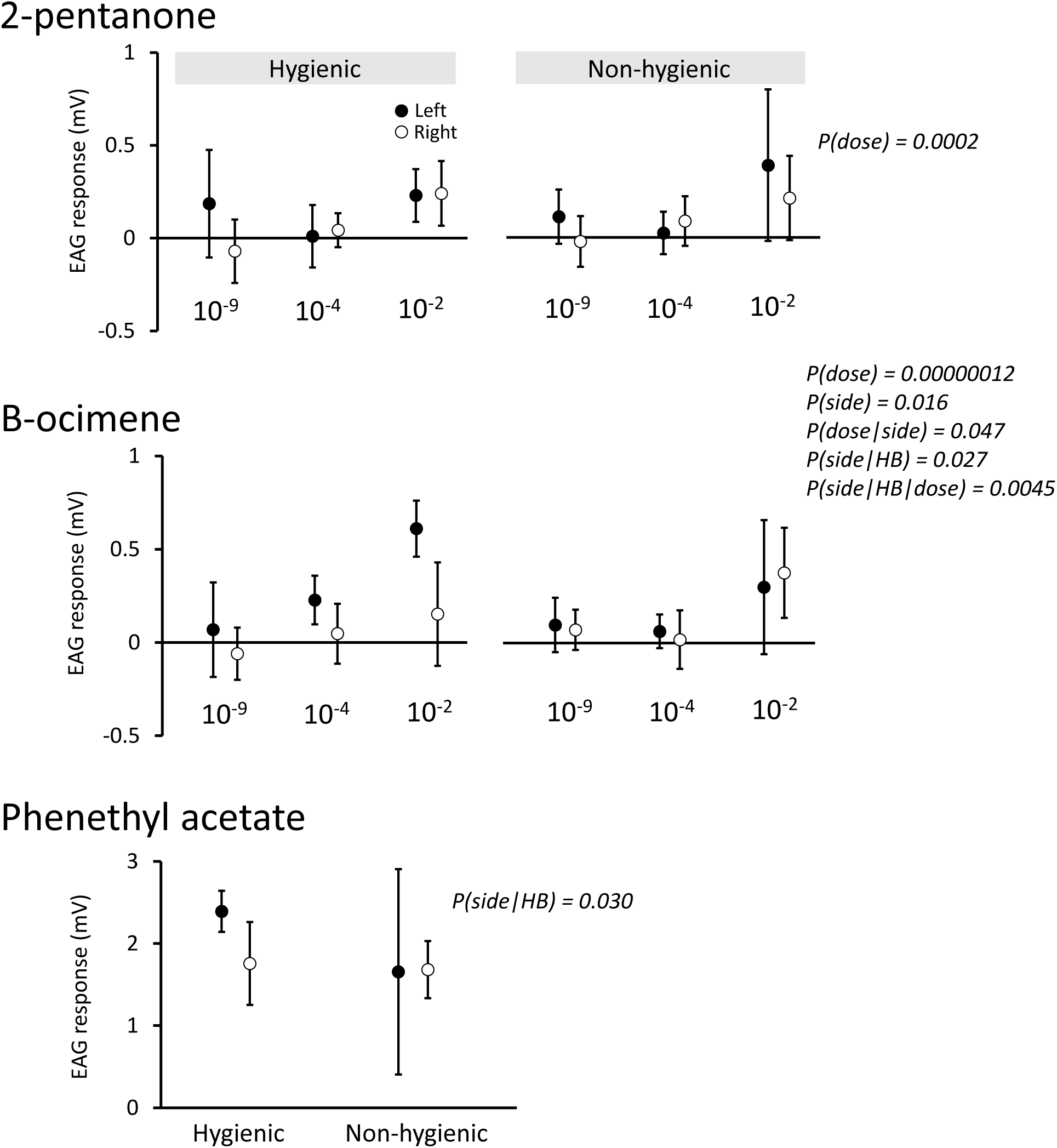
Antennae nerve stimulations of candidate HB-inducing compounds. Electroantennography was used to quantify antenna nerve responses to odour stimuli. 2-pentanone and β-ocimene doses were applied at three dilutions (10_-9_, 10_-4_ and 10^−2^ v/v; N = 5-9). Phenethyl acetate, a known HB-inducing compound, was applied at one dose (10^−9^ v/v).

A well-known phenomenon in olfactory perception is the synergistic effect of odorant mixtures^37^. That is, mixtures can be perceived not as the sum of their parts, but as if they are entirely new odours; however, this is rarely observed in honey bees^38-40^. To test if the four odours could lead to stronger EAG signals by stimulating antennae synergistically, we produced equivolume mixtures (1% total in ethanol) of all possible combinations of isopropanol, 2-pentanone, β-ocimene and oleic acid and used these to perform EAG on left antennae of hygienic bees. As expected, none of the odour combinations induced greater antenna stimulations than β-ocimene alone (the strongest stimulator; **Appendix Figure S2**).

### No proteomic differences were observed between left and right antennae

To determine a potential mechanism for lateralization of antenna stimulation at the gene expression level, we performed quantitative proteomics on left and right antennae of nurse bees from five hygienic colonies. Despite identifying 1,845 proteins (13,128 peptides), none of them were differentially expressed (**Appendix Figure S3**). Interestingly, 230 of the identified proteins are ones that were discarded from the first Official Gene Set (OGSv1.0), apparently in error. We described this phenomenon previously^41^ and this finding offers secondary confirmation. A further 15 proteins are new sequences which we identified in the same previous proteogenomic effort.

## Discussion

Overall, our experimental findings point to emerging mechanistic patterns regarding HB. We found that a well-known brood pheromone, β ocimene, was strongly emitted from FKB and this pattern positively correlates with HB score. We also identified one compound, oleic acid, which was consistently released in higher amounts in FKB, not only across colonies but also across developmental stages. Finally, we functionally validated these compounds using electroantennography and show that lateralization of antennal response is strongly associated with HB, but we could not identify associated proteomic changes. Unlike the FKB, we found that *Varroa*-infestation causes subtle but significant changes to the overall cuticle compound profile, although no individual compounds emerged as drivers. This may in part explain why trait selection for *Varroa*-sensitive hygiene, a specialized form of HB, requires more rigorous selection techniques.

The clear majority of differentially emitted compounds were more abundant in dead pupae than in live ones. This is intuitive, since HB-triggering compounds should give a more reliable and specific signal to the bees if dead:live discrimination is based on their presence, rather than absence. Interestingly, three different likely terpene peaks were identified (based on a MS2 base peak of 93.0 m/z and spectral matches to other terpenes; Table 1) but of these, only β-ocimene could be confidently confirmed. These unidentified compounds could still certainly be biologically relevant to HB, but further work is required to identify them.

Oleic acid – an omega-9 monounsaturated fatty acid – was emitted more strongly in FKB compared to live brood of all ages tested in this study. Intriguingly, oleic acid has been implicated as a mechanistic agent for HB in other social insects^42,43^. For example, Wilson *et al.*^43^ found that applying oleic acid to otherwise mobile and healthy ants induced other ants to transport them to their refuse area. Oleic acid also binds strongly to odorant binding protein 18, which significantly correlates with HB in honey bees^16^ and is currently being used for marker-assisted selective HB breeding^17^. Finally, it has been shown previously that *Varroa*-parasitized brood (which can also trigger HB) emits more oleic acid compared to healthy brood; however, at the time of that study it was not identified as a discriminating compound^30^, and we could not replicate these results in our analyses of *Varroa*-infested pupae. Oleic acid did not elicit a strong EAG signal, however, possibly because of its low volatility (oleic acid boiling point: 360°C; 2-pentanone was the next highest at 101°C).

β-ocimene is a brood pheromone^44-46^ normally released by young larvae in order to stimulate adult workers to feed them^34^, but, to our knowledge, its release has not been previously associated with the FKB assay or HB. It seems unusual that dead bees would emit more of a brood pheromone than live bees, but it could be that a normally tightly controlled pheromone release mechanism breaks down as membranes become more permeable after freezing. β-ocimene was also the tallest peak in the chromatograms and was the only compound to elicit both dose-dependent and HB-dependent EAG responses (along with the known HB-inducing compound: phenethyl acetate; Fig. 5). This finding is intriguing for two reasons: 1) β-ocimene has previously been shown to increase the frequency of worker visits to brood^34^ and 2) an independent study found that a different brood pheromone (brood ester pheromone; BEP) was also significantly more abundant in parasitized brood^33^. By increasing worker visits to brood cells that should otherwise not require attendance, β-ocimene may attract the attention of bees that can perform HB. As the second brood pheromone implicated in HB, these results may indicate a broader pattern of HB dependence on brood pheromones. Furthermore, Mondet *et al.*^33^ suggest that BEP contributes to detection of *Varroa*-infested pupae by signaling developmental delay – this mechanism is consistent with our own observations, since more β-ocimene is emitted from larvae compared to pupae^44^.

Further stimulating our interest in this compound, we also found that the ratio of β-ocimene emitted from FKB to live pupae significantly positively correlates with HB itself. This suggests that there may be a brood effect contributing to HB scores, in addition to olfactory sensitivity of adult workers, even though previously this was not thought to be the case. In an early foundational paper, Spivak and Downey^20^ found no brood effect when they performed hygienic tests using reciprocally donated brood; however, brood age was not controlled during these tests. In the same study, they established that brood age had a significant effect on non-hygienic colonies but not on hygienic colonies, with non-hygienic colonies performing significantly better on the FKB test when young brood (capped larvae and prepupae) was used compared to older pupae. The confounding factors may have simply erased potential brood effect patterns. Interestingly, β-ocimene is also present in far higher amounts in larvae compared to pupae and the larva cuticle is more delicate, so it is possible that disruption of the larval membranes by freeze-killing leads to an even larger proportion of β-ocimene sample size (n = 8) does not accurately represent all colonies and our work could likely benefit from a larger scale experiment.

This is not the first time that a brood effect has been suggested: Parker *et al.* (2012) found significant differences in the larval cuticle proteome between high and low VSH bees (a specialized form of HB targeting *Varroa* mites) and suggested that this may lead the brood to emit different chemical cues. This data, together with our own, suggests that HB could be dependent on two interacting factors – the strength of brood odour and the workers’ limit of odour detection – rather than the adult workers’ olfactory sensitivity alone.

The left-biased EAG response lateralization is intriguing. Lateralization in bees is not new: Rogers *et al.*^47^ have shown that bees are more likely to interact aggressively when communicating *via* their left antenna, whereas they have preferentially positive encounters when interacting *via* their right antenna. Interestingly, Rogers and Vallortigara^48^ found that bees performed better at long term memory recall tasks when stimulated *via* their left antennae, but not their right. We did not acquire data on the higher order processing of odours, but these studies create a precedent for antenna lateralization as it relates to behaviour. It is curious that despite having more olfactory sensilla on the right antenna^35^, the left elicits a stronger EAG signal for FKB and chalkbrood compounds. One possible explanation is that the olfactory sensilla that do exist on the left antenna house olfactory receptor neurons that are specifically tuned to particular odours.

Given the apparent lateralization of response observed here and electron micrographs showing more olfactory sensilla on the right antennae of worker bees^35^, we expected to also find differences in protein expression between left and right antennae of hygienic bees (**Appendix Figure S3**). The lack of any detectable difference indicates that we either did not penetrate deep enough into the proteome or that the contribution of differential expression in different sensilla is small compared to the overall expression of the relevant proteins. In the future, our proteomics analysis could be improved by performing sample fractionation to increase depth.

When we compared odour profiles of *Varroa*-parasitized pupae to healthy pupae across four developmental stages, we found a significant interaction between parasitization and developmental stage but no individual compounds drove this effect (Fig 4). This could be because VSH is a specialized form of HB^49^ and this specialization is required because the differences between infested and non-infested brood are subtler than for dead and live brood. Indeed, one strategy for mites to evade detection in the colony is to adapt its own cuticle hydrocarbon profile to mimic its host^50^. Another explanation could be that since the healthy control brood was pulled from cells immediately adjacent to the infested pupa, it could be that *Varroa*-associated compounds transferred through the thin wax wall to the healthy pupae, diminishing the observable differences. However, we still believe that this was the appropriate comparison, since hygienic bees must be able to discriminate between the healthy and diseased states. Finally, it could also be that key differentially emitted compounds do exist, but we were unable to detect them with our methods. Mondet *et al.*^33^ were recently able to find *Varroa*-specific compounds by analyzing solvent extracts of crushed infested pupae, although it is not clear that compounds measured in this way would be detectable by bees performing HB. The hexane extraction and SPME used here are suitable for capturing non-polar compounds with relatively high volatility but it could be that the superior olfactory sensitivity of hygienic bees allows them to detect some polar, non-volatile compounds. Indeed, oleic acid (a carboxylic acid) is one of our most confident HB-inducing candidates but it was among the last to elute in our GC-MS analysis of hexane extracts; more polar compounds would likely become trapped in the GC-MS inlet or not be miscible in hexane at all.

## Conclusion

The work presented here furthers our understanding of HB and the underlying mechanism. Interestingly, this is now the second study to implicate a known brood pheromone (β-ocimene, in this case) as a mechanistic agent for HB. We found that hygienic bees, but not non-hygienic bees, elicit a dose-dependent lateralized olfactory response to β-ocimene. Furthermore, it is already known that odorant binding protein 18, which is thought to aid in odour detection, positively correlates with HB and here we show that one of its strongest known ligands (oleic acid) is indeed abundantly emitted from dead brood. This compound is also known to induce HB in other social insects, suggesting that the mechanism for HB is evolutionarily conserved. The odour profiles of *Varroa*-infested brood showed a significant interaction between infestation and developmental stage, and this subtlety is consistent with VSH being a specialized form of HB. Further experiments are needed to confirm the identities of the five unknown significant compounds since they may still be biologically relevant.

## Materials and methods

### Honey bee colonies and hygienic testing

Honey bee colonies were kept at three separate locations in Greater Vancouver, BC, Canada. Colonies were scored for HB using the FKB assay as previously described^20^. All testing and sampling was conducted during the summer of 2016.

### FKB GC-MS sample collection

Honey bee pupae with no visible signs of disease were collected from colonies by carefully uncapping cells and removing pupae with clean stainless steel forceps. Age was determined based on eye and cuticle pigment using the following relationships: white-eyed = 12-13 d, pink-eyed = 14-15 d, purple-eyed white body = 16 d and purple-eyed tan body = 17-18 d. From bee to bee, eye and cuticle pigment was matched exactly so that each bee in each age group was at the same developmental stage. Pupae were placed in clean glass vials, removing any wax debris and avoiding abrasions or cuticle indentations. Freeze-killed samples were placed at −80°C (15 min) then placed in a humid 33°C incubator (24 h), while live samples were placed directly into the same incubator.

Compounds were extracted for low resolution GC-MS analysis by two different methods: solvent extraction and solid phase micro-extraction (SPME). For analyzing cuticular compounds across developmental stages (white-eyed, pink-eyed, purple-eyed white body and purple-eyed tan body; n = 3), extracts were prepared by washing whole pupae with 300 µl HPLC-grade hexane for 5 min with gentle agitation. Hexane extracts were transferred to a clean vial and immediately stored at −80°C until GC-MS analysis. For the cross-colony analysis (N = 3 per colony, n = 6 colonies), compounds were extracted only from purple-eyed white body pupae using the method above as well as by sealing individual freeze-killed and live pupae in 10 mL glass vials (Supelco) and incubating at 33°C (24 h) for SPME analysis. We confirmed that 10 mL of air is enough for one bee to survive for this time by performing the same procedure for late-stage pupae, which were still actively moving after being sealed for 24 h.

One µL of each hexane extract was analyzed by GC-MS (Agilent 6890N/5975C Inert XL MSD) using a DB-wax column (J&W 122-7032) and a 30 min gradient from 50°C to 230°C. The back inlet (pulsed splitless) was at 250°C and 6.24 psi with a 53.5 mL/min flow rate (He gas) connected to the analytical column (30 m, 250 µm ID). The instrument was set to scan from 40 – 300 m/z. The MS source and quadrupole were maintained at 230°C and 150°C, respectively.

Headspace volatiles were sampled using solid phase micro-extraction (SPME) and analyzed by GC-MS (Agilent 7890A/5975C Inert XL MSD) using a 45 min gradient and the same column model as above. We used a 50/30 µm DVB/CAR/PDMS stableflex SPME fiber and sampling details were: 40°C incubation, 3 s agitation at 500 rpm, 600 s extraction time and 300 s desorption time. The oven settings were: 35°C (stable; 4 min), then 25°C/min (5 min) and a 2:1 split ratio. The inlet temperature was 250°C and MS acquisition parameters were the same as above except that the lower mass limit was 33 m/z.

### Varroa destructor GC-MS sample collection

For ease of sampling, mite-infested brood was concentrated on a single frame by caging the queen in a single-frame excluder and transplanting all other open brood into a temporary ‘incubator’ colony. This left only the single frame of brood suitable for mite infestation, effectively concentrating the phoretic mites looking for brood cells in that colony to one location. After 10 d, the brood was returned from the incubator colony and the queen was released. Following this, pupae were sampled by the same methods as above and only pupae with a single foundress mite were chosen. The accompanying mite family (including foundresses, deutonymphs, protonymphs and eggs; Fig. 1) was transferred to the same glass vial as the pupa using a soft paintbrush. Adjacent, age-matched non-infested sister pupae with no visible signs of disease were collected as controls.

### GC-MS data analysis

GC-MS data was analyzed using Mass Hunter Qualitative Analysis software (vB.06.00). Chromatogram peaks were first smoothed using the default algorithm and then manually integrated to ensure consistent baselines between replicates. To compare FKB to healthy odour profiles across developmental stages, peak areas were exported to Excel (2013) where they were log_10_ transformed and groups (developmental stage, treatment, compound type) were compared using three-factor ANOVA (Excel), followed by a Tukey HSD *post-hoc* test to identify the specific differentially emitted compounds. We did not test the data for normality, but the ANOVA is generally tolerant to non-normal data and/or low replication. The same process was used to analyze FKB changes across colonies except that a two-factor ANOVA was employed since this only involved a single developmental stage (purple-eyed white body pupae). To determine if any significantly differentially emitted compounds correlated with colony HB score, we calculated the dead:live ratio (not log transformed), then the Pearson correlation for each one. Significance was determined by comparing P-values against the Bonferonni-corrected α. The effect of *Varroa*-infestation was also examined using a three-factor ANOVA (developmental stage, infestation, compound). In all cases, compound identities were determined by searching spectra against the Wiley Chemical Compound Library (W9N08.L) in Mass Hunter.

### Antenna preparation for electroantennography

Bees for electroantennography (EAG) were collected across two colonies with high HB scores and three with low HB scores. Since bees perform HB best when they are two to three weeks old^51^ we marked emerging bees with a paint pen and returned them to the hive for 14 days, then EAG data was acquired for up to one week. Antennae were excised and both ends were trimmed with a scalpel, randomizing whether right or left antennae were excised first. Trimmed antennae were then attached to glass capillary reference and recording electrodes filled with insect saline solution (210mM NaCl, 3.1mM KCl, 10mM CaCl_2_, 2.1mM NaCO_3_, 0.1 NaH_2_PO_4_) as previously described^52^. EAG responses were recorded on the EAD program of a Syntech™ IDAC-4 signal acquisition unit. The low cutoff was set at 0.1 Hz, high cutoff at 10 Hz, external amplifier set to 1. Humidified, charcoal filtered air was passed continuously over the antenna via a Syntech CS-55 stimulus controller, also serving as a carrier for odour-filled pulses. Odorants were dispensed onto 1 cm^2^ No. 1 Whatman filter paper, allowing the solvent to evaporate for 30 s before being inserted into a glass Pasteur pipette. Odorant pulses were passed through the Pasteur pipette to the antenna for 1 s, and 0.5-1 minute was allowed between each presentation of an odour for the antenna to return to baseline activity. Each antenna was stimulated with a series of three dilutions (10^−9^, 10^−4^ and 10^−2^ v/v in ethanol) each of isopropanol, 2-pentanone, β-ocimene and oleic acid (all from Sigma or Fisher; >90% purity). Phenethyl acetate, a known HB-inducing compound isolated from chalkbrood^25^, was used as a positive control. All possible equivolume combinations of the four candidate compounds were also tested at a 10^−2^ (1%) dilution to test for synergistic effects of mixtures.

Even though antennae were conditioned with humidified air throughout the recordings, EAG signal decay was still evident even for stimuli of solvent alone over time. Therefore, each antenna was subject to intermittent solvent stimulations throughout the recordings to mathematically interpolate the background solvent stimulus. The quality cut-off for the solvent curve fit was R^2^ > 0.9: traces which did not meet this criterion were discarded. Since the number of surviving traces varied, in total we acquired between 5 and 9 biological replicates in each experimental group (left vs. right; high HB vs. low HB). Finally, the interpolated solvent amplitude was then subtracted from the solvent + odour stimulations, resulting in the mV value that can be attributed to the odour alone. The amplitudes of our recordings are consistent with other similar studies in bees^22,24,48^. Statistical analyses were conducted in Excel.

### Antenna protein extraction and proteomic analysis

Thirty to forty bees on open brood frames were collected from five highly hygienic colonies (n = 5; all with FKB scores > 94%; Table 2). Bees were anesthetized with carbon dioxide and their antennae dissected on ice followed by homogenization (Precellys 24; Bertin instruments) with ceramic beads (lysis buffer: 6M guanidinium chloride with 10 mM TCEP, 100 mM Tris (pH 8.5), 40 mM chloroacetamide). The homogenizer was set to 6,400 M/s for 30 s × 3 (1 min on ice in between). Lysate was transferred to a new tube and debris was pelleted (16,000 rpm, 15 min, 4°C), followed by acetone precipitation as previously described^53^. Dried protein pellets were resuspended in 50 mM ammonium bicarbonate buffer (1% sodium deoxycholate) and protein concentration was determined using a bicinchoninic assay (Pierce). Protein was reduced, alkylated, digested and analyzed on an LC-ESI-MSMS system (Easy nLC-1000 coupled to a Bruker Impact II mass spectrometer) as described in our previous publication^41^, except we loaded 2.5 µg (based on protein quantitation), the LC gradient was 165 min and MS/MS frequency was set to 18 Hz (see embedded microTOFQImpactAcquisition.method files within PXD005242 for further details).

**Table 2.**
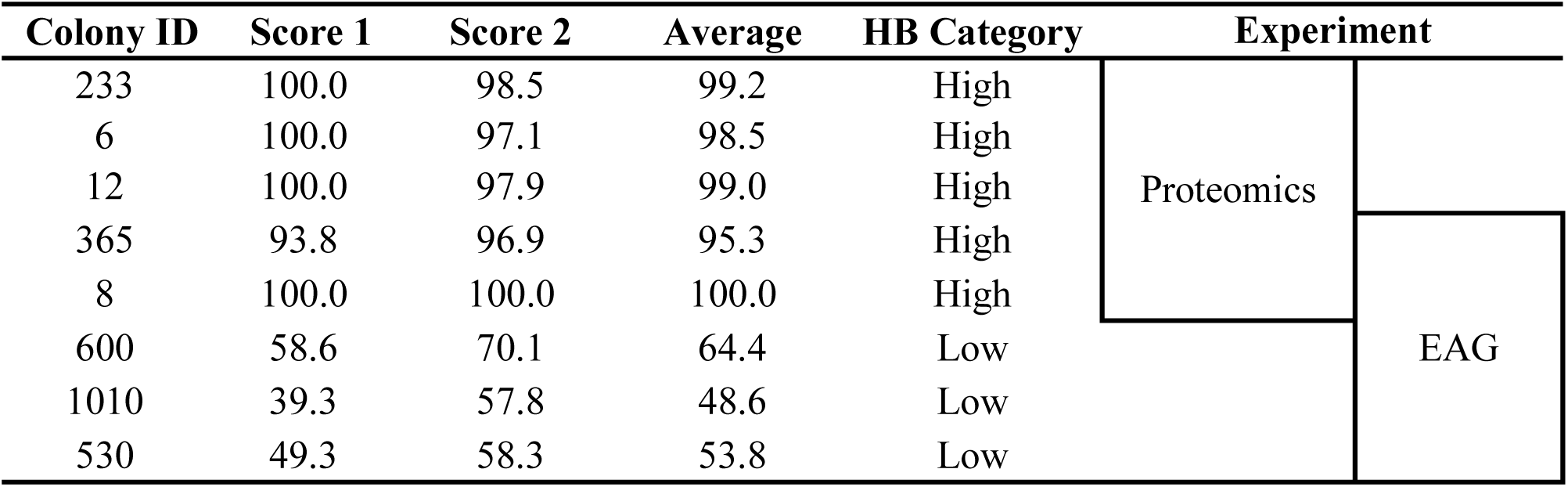
FKB scores

Proteomics data was searched using MaxQuant (v1.5.5.30) and processed using Perseus (v1.5.5.3). All data was deposited to ProteomeXchange (PXD005242). All MaxQuant search parameters were left as default except: deamidation (NQ) was added as a variable modification, “match between runs,” “label-free quantification” and “re-quantification” options were enabled and “min ratio count” was set to 1. Briefly, reverse hits, proteins “only identified by site” and contaminants were removed followed by filtering for proteins identified in four or more colonies. Data was then Log2 transformed and missing values were imputed before comparing left and right antennae using a two-group comparison in Perseus.

## Acknowledgements

We would like to acknowledge Dr. Joerg Bohlmann for allowing us to use the GC-MS facilities and providing samples of synthetic chemical standards for confirming compound identifications.

AM was supported by a Natural Sciences and Engineering Research Council Canada Graduate Scholarship Doctoral award (NSERC CGS-D). The work described here was funded by an NSERC Discovery Grant to LJF.

## Author contributions

AM designed most of the experiments, with assistance from LJF and LLM. TFC performed the EAG experiments. LLM ran the GC-MS samples and provided GC-MS consultation. AM wrote the first draft of the manuscript and prepared the proteomics samples. AM collected most samples and analyzed most of the data, with some assistance from TFC.

